# Factors required for adhesion of *Salmonella enterica* serovar Typhimurium to lettuce (*Lactuca sativa*)

**DOI:** 10.1101/2020.04.03.024968

**Authors:** Laura Elpers, Michael Hensel

**Affiliations:** Abt. Mikrobiologie, Universität Osnabrück, Osnabrück, Germany; CellNanOs – Center for Cellular Nanoanalytics, Universität Osnabrück, Osnabrück, Germany

**Author notes:** Address for correspondence: Michael Hensel, Abteilung Mikrobiologie, CellNanOs – Center for Cellular Nanoanalytics Osnabrück, Fachbereich Biologie/Chemie, Universität Osnabrück, Barbarastr. 11, 49076 Osnabrück, Germany.

**Keywords:** adhesiome, *Salmonella enterica* serovar Typhimurium, lettuce, *Lactuca sativa*, Lpf fimbriae, Sti fimbriae, MisL, BapA, LPS, flagella-mediated motility

## Abstract

*Salmonella enterica* serovar Typhimurium (STM) is a major cause of food-borne gastroenteritis. Recent outbreaks of infections by STM are often associated with non-animal related food, i.e. vegetables, fruits, herbs, sprouts and nuts. One main problem related to consumption of fresh produce is the minimal processing, especially for leafy salads such as corn salad, rocket salad, or lettuce. In this study, we focused on lettuce (*Lactuca sativa*) which is contaminated by STM at higher rates compared to corn salad, resulting in prolonged persistence. We previously described the contribution of Saf fimbriae, type 1 secretion system (T1SS)-secreted BapA, intact LPS, and flagella-mediated motility to adhesion to corn salad leaves. We systematically analyzed factors contributing to adhesion of STM to lettuce leaves. We used the previously established reductionist, synthetic approach to identify factors that contribute to the surface binding of STM to leaves of lettuce by expressing all known adhesive structure by the Tet-on system. The analyses revealed contributions of Lpf fimbriae, Sti fimbriae, autotransported adhesin MisL, T1SS-secreted BapA, intact LPS, and flagella-mediated motility to adhesion of STM to lettuce leaves. In addition, we identified BapA is a potential adhesin involved in binding to corn salad and lettuce leaf surfaces.

**Importance:** Gastrointestinal pathogens can be transmitted by animal products, as well as by fresh produce of non-animal origin. The numbers of outbreaks by fresh produce contaminated with gastrointestinal pathogens are increasing, and underline the relevance to human health. The mechanisms involved in the colonization of, persistence on, and transmission by fresh produce are poorly understood and have to be part of further research. Here, we investigated the contribution of adhesive factors of STM in the initial phase of plant colonization, i.e. the binding to the plant surface. Usage of a reductionist, synthetic approach including the controlled surface expression of specific adhesive structures of STM, one at a time, allowed the determination of relevant fimbrial and non-fimbrial adhesins, the O-antigen of lipopolysaccharide, the flagella, and chemotaxis to binding to lettuce leaves.

## Introduction

*Salmonella enterica* is a common cause of food-borne gastroenteritis leading to thousands of fatal cases worldwide (1). Currently, the numbers of outbreaks of *S. enterica* infections associated with consumption of fresh produce are increasing (2). Besides contaminated products of animal origin, i.e. eggs, chicken or pork, fresh produce like vegetables, fruits, salads, herbs, and nuts may be contaminated by *S. enterica* (3). Contamination can occur during pre-harvest by the seeds themselves, by irrigation water, or by fertilizers (often based on animal source). After harvest, fresh produce may be contaminated by improper implemented hygiene regulations, or additional processing steps (washing water, cutting). After first attachment and adhesion, *S. enterica* can colonize and persist in and on plants, leading to possible replication and further spreading (4). Several studies investigated the interaction of *S. enterica* and various salad species, thereby focusing on the important first contact of *S. enterica* to salad. It was shown that *S. enterica* serovar Thompson adheres to spinach leaves in higher numbers compared to watercress leaves. Attachment to abaxial and adaxial leaf surfaces was comparable (5). Differences in attachment levels to various salad species were further investigated for S. en*terica* serovar Typhimurium (STM). Adhesion of STM to lettuce leaves occurred in higher numbers compared to corn salad leaves, moreover persistence *in planta* was longer for lettuce than for corn salad (6).

For the adhesion of *S. enterica* to salad, different factors have been investigated, i.e. involvement of flagella. Directed motility is involved in adhesion to iceberg lettuce leaves while loss of flagella filaments ablated binding of STM. The binding of a smooth swimming Δ*cheY* strain was not affected, whereas the internalization was prevented (7). Other studies addressed the involvement of genes in long-term persistence on lettuce leaves under cold storage. Work of Y. Kroupitski et al. (8) revealed a possible involvement of *bcsA, misL* and *yidR* due to decreased attachment to lettuce leaves and decreased survival during cold storage for respective mutant strains.

Investigations of *Salmonella*-plant interactions are complex, since different sites of contamination (roots, leaves, fruits, seeds), age of leaves, and possible also roots, fruits and seeds, metabolic changes of plants during dark-night rhythm, varying temperatures, and other environmental stresses (e.g. UV light) have to be considered. Here, we focus on the adhesion of STM to lettuce leaves, and aim to reveal factors involved in initial colonization and *Salmonella*-plant interaction. While previous studies mainly investigated STM WT, or strains mutated in candidate adhesion factors, we focus on expression of defined adhesive structures, one at a time, to reveal possible ligands on lettuce leave surfaces.

STM possesses a complex set of adhesive structures including 12 chaperone-usher (C/U) fimbriae, curli fimbriae assembled by the nucleation-precipitation pathway, two type 1 secretion system (T1SS)-secreted adhesins (BapA, SiiE), three type 5 secretion system (T5SS)-secreted (autotransported) adhesins (MisL, ShdA, SadA), and two outer membrane proteins (OMPs) with putative adhesive features (PagN and Rck) (9, 10). For most C/U fimbriae, little is known about conditions of native expression and binding properties (11). All operons encoding C/U fimbriae consists at least of a fimbrial main subunit, a specific periplasmic chaperone, and an usher located in the outer membrane (12). Best characterized C/U fimbriae so far are the mannose-sensitive type 1 fimbriae encoded by *fimAICDHF* (13), and *pef* fimbriae encoded by genes on pSLT2. Pef fimbriae are involved in the binding to Le^x^ blood group antigen and in the formation of biofilm on chicken intestinal epithelium (14, 15). Curli fimbriae are encoded by two operons *csgBAC* and *csgDEFG*, and are assembled by the nucleation-precipitation pathway. Curli fimbriae are known to be involved in the formation of biofilms together with cellulose (16, 17).

The *Salmonella* pathogenicity island 4 (SPI4) locus (*siiABCDEF*) encodes the T1SS (SiiCDF) which mediates the secretion and surface expression of giant adhesin SiiE on the bacterial surface (18). SiiE consists of 53 repetitive bacterial Ig domains (BIg) and is the largest protein in STM with a molecular mass of 595 kDa. SiiE specifically binds glycostructures with N-acetylglucosamine (GlcNAc) or 2,3-linked sialic acid (19). SiiE mediates the first contact of STM to polarized epithelial cells, which is followed by invasion mediated by the SPI1-encoded T3SS and various effector proteins (20, 21). The other T1SS-secreted adhesin BapA (386 kDa) is encoded by *bapABCD* (<u>b</u>iofilm <u>a</u>ssociated <u>p</u>rotein), contains 28 BIg domains, and is involved in biofilm formation (17, 22). The T1SS for surface expression of BapA is composed of BapBCD. STM possesses two monomeric autotransporters (MisL and ShdA) and one trimeric autotransporter (SadA). These autotransporters are not expressed under laboratory conditions and little is known about their native expression (23). The monomeric autotransporters MisL and ShdA are known to bind fibronectin, thus being involved in the intestinal infection of mice (24-26). SadA of STM strain SL1344 is possibly involved in biofilm formation and adhesion to CaCo2 cells but only in a strain with altered lipopolysaccharide (LPS) (27). In addition to adhesins proper, as possible factors involved in adhesion to lettuce leaves LPS, flagella filament and motility were considered.

For most adhesins, general environmental or host factors inducing expression are not known. Therefore, we used a synthetic system based on the *tetA* promoter for controlled expression of various STM adhesins (23). We tested the adhesive structures for involvement in binding to lettuce leave surfaces by a reductionist approach described before (28). Our data will support the classification of adhesive structures commonly involved in binding to salad species, in order to subsequently develop strategies for prevention or reduction of adhesion of *S. enterica* to salad leaves.

## Results

In a previous study (28) we deployed a reductionist, synthetic approach to identify factors that contribute to the surface binding of STM to leaves of corn salad. We now investigate factors that contribute to surface binding of STM to leaves of lettuce. Infection of corn salad and lettuce grown under aseptic conditions with STM indeed revealed a higher adhesion of STM to lettuce (Figure S1B). To identify adhesive structures required for adhesion to multiple or individual salad species, we investigated the complex adhesiome of STM in its entirety.

Various databases such as GEO, SalCom, and others (6, 29, 30) were used to identify potential environmental stimuli that induce expression of STM adhesins. However, these analyses only revealed defined culture conditions leading to expression of 3 (i.e. Saf, Sii, PagN) of 20 adhesins in the STM adhesiome (Janina Noster, MH, data not shown). The rather diverse nature of these inducing conditions excluded a systemic comparison. Therefore, we expressed various adhesins ectopically under control of the *tetR* P_*tetA*_ cassette as previously described (23). To avoid interference with native expression of certain adhesins, we decided to use strain SR11 Δ12 lacking all 12 C/U fimbriae. STM SR11 Δ12 strains harboring plasmids for expression of various adhesins were tested for contribution to adhesion to lettuce by validating the level of adhesion. Further, we tested strains lacking further putative adhesion factors. These strains included defects in flagella assembly and motility, in LPS structure, and a strain lacking all 20 known adhesive structures (Δ12 Δ*misL* Δ*sadA* Δ*shdA* ΔSPI4 Δ*bapABCD* Δ*rck* Δ*pagN* Δ*csgBAC-DEFG* = SR11 Δ20). Adhesion assays with lettuce leaves revealed no differences in adhesion for SR11 WT and the used background SR11 Δ12 lacking all 12 C/U fimbriae (Figure S1A). Furthermore, the deletion of all putative adhesive structures, strains SR11 Δ20 was not altered in adhesion compared to SR11 Δ12 (Figure S1C).

### Contribution of fimbrial adhesins to adhesion to lettuce

We analyzed adhesion to lettuce by anhydrotetracycline (AHT)-induced expression of all C/U fimbriae encoded by STM. Different levels of binding to lettuce leaves were mediated by expression of C/U fimbriae (Figure 2A). Expression of Bcf, Sth, Pef, Stb and Stj did not affect adhesion compared to background strain SR11 Δ12. Therefore, cognate ligands may be absent on lettuce leaves. The expression of Fim, Saf, Stc, Std and Stf fimbriae led to decreased adhesion (47%, 73%, 59%, 76%, and 72% mean, respectively). However, SR11 Δ12 harboring plasmids encoding for Saf, Stc and Stf fimbriae also resulted in decreased adhesion in the absence of inducer AHT. In absence of AHT no synthesis and transport to bacterial surface of these three fimbriae were observed by flow cytometry (23). Expression of Lpf and Sti fimbriae resulted in significantly increased adhesion to lettuce (means of 167% and 139%, respectively). Non-induced controls for Lpf and Sti fimbriae showed similar adhesion compared to background SR11 Δ12. Hence, we propose cognate ligands on lettuce leaves for Lpf and Sti fimbriae. AHT-induced expression of curli fimbriae led did not alter adhesion (Figure 2***Error! Reference source not found.***B), whereas the deletion of *csgBAC* and *csgDEFG* resulted a significantly increased adhesion to lettuce (131% mean).

**Figure 1:**
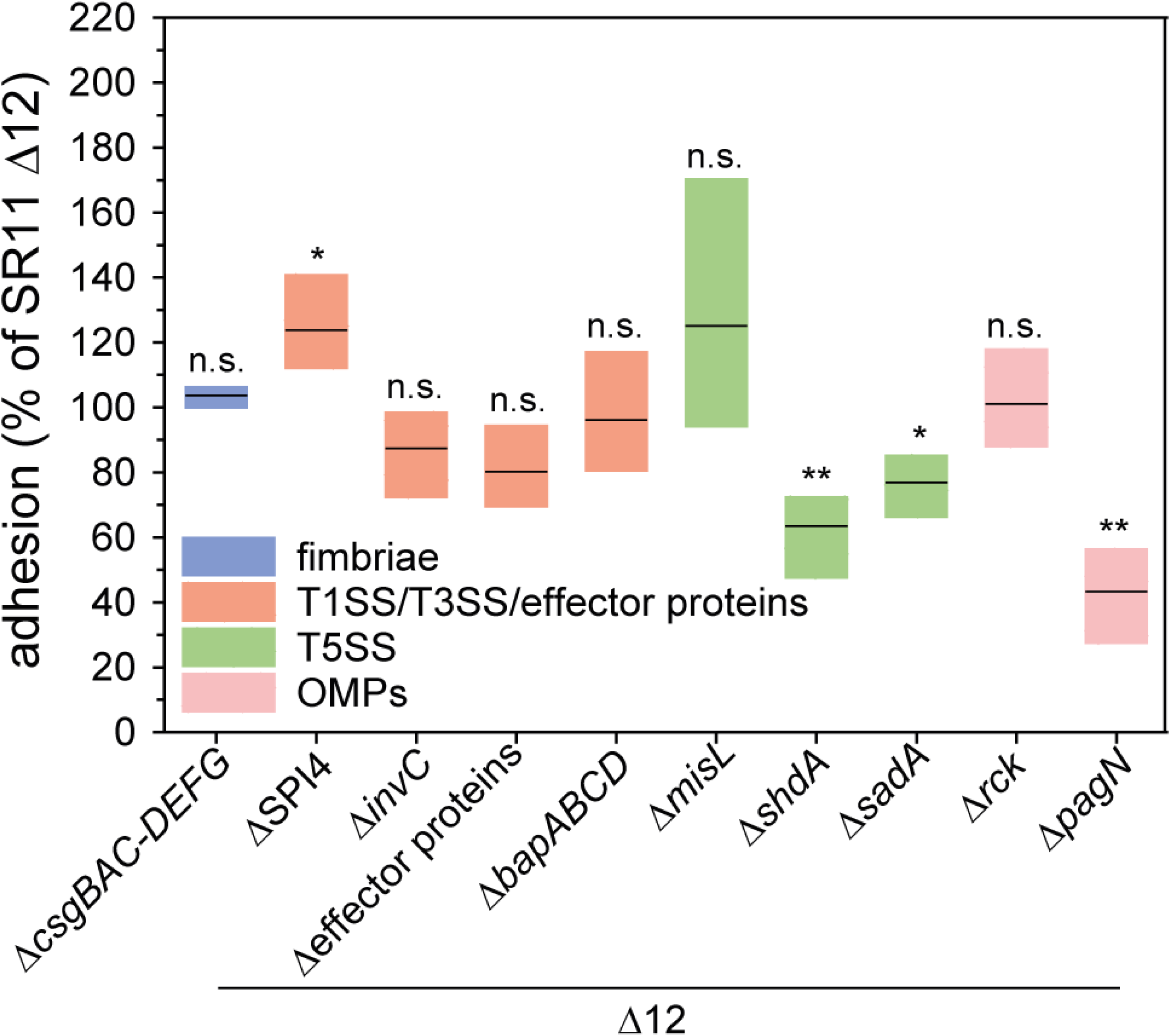
Impact of genes encoding putative adhesive structures and effector proteins of SPI1-T3SS on adhesion of STM to lettuce. Lettuce was grown under aseptic conditions, leave disks were generated, and infected with STM SR11 Δ12, or various SR11 Δ12 strains with additional deletions in genes encoding putative adhesive structures, and effector proteins of the SPI1-T3SS (Δ*sopA* Δ*sopB* Δ*sopD* Δ*sopE2* Δ*sipA* = Δeffector proteins). Overnight cultures were diluted 1:31 in fresh LB, bacteria were subcultured for 3.5 h and diluted in PBS for infection of lettuce leave disks. After infection for 1 h, lettuce leave disks were washed three times to remove non-adherent bacteria. For the quantification of adherent bacteria, leave disks were homogenized in PBS containing 1% deoxycholate, and serial dilutions of the homogenate and inoculum were plated onto MH agar plates for the quantification of CFU. Levels of adhesion were determined as percentage of inoculum CFU recovered in leave disk homogenates, and adhesion of various strains was normalized SR11 Δ12 set to 100% adhesion. The distributions of three biological replicates is represented as box plots with means and median values. Statistical significances were calculated by the Student’s *t* test and are indicated as follows: n.s., not significant; *, *P* < 0.05; **, *P* < 0.01; ***, *P* < 0.001.

**Figure 2:**
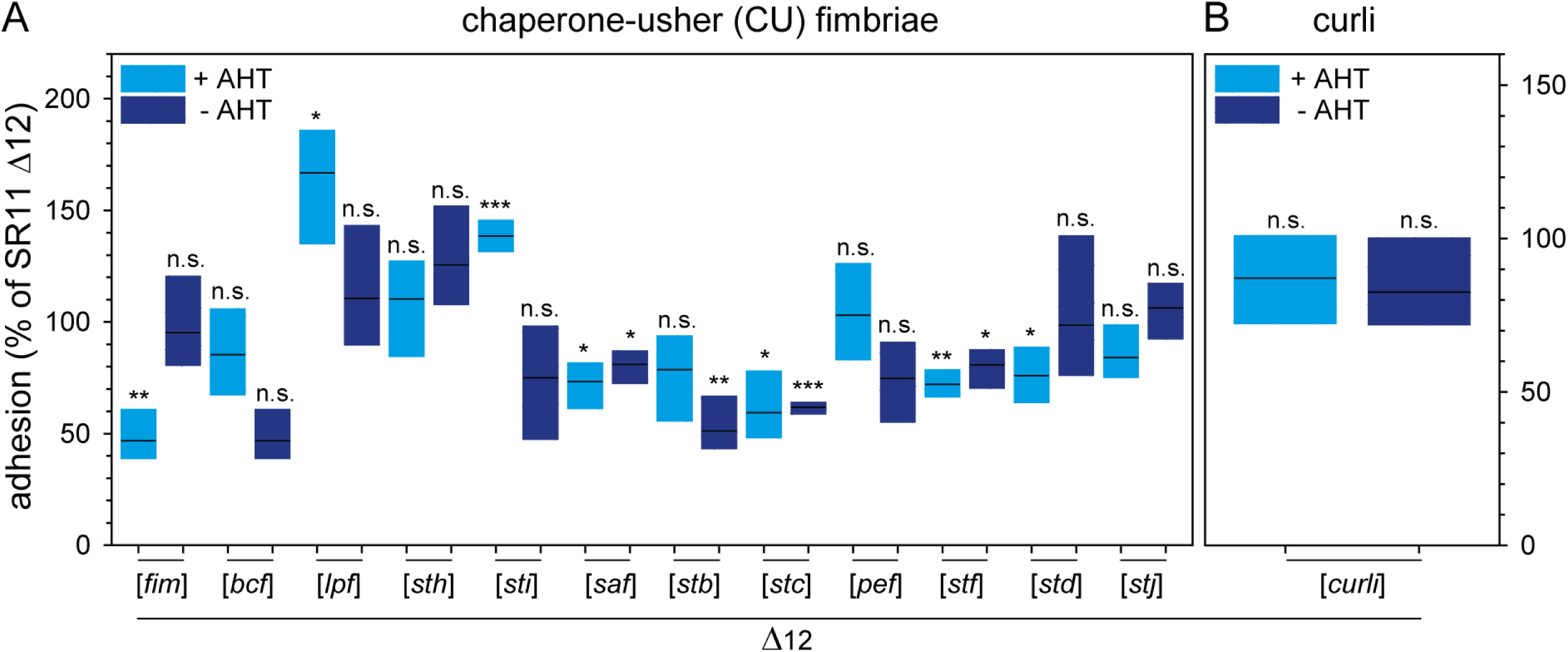
Impact of chaperone-usher fimbriae and curli fimbriae expression on adhesion of STM to lettuce. Sterile grown lettuce was infected with STM SR11 Δ12, of STM SR11 Δ12 strains harboring plasmids for the expression of various chaperone-usher fimbriae (A), or curli fimbriae (B). If indicated, expression of fimbriae was induced by addition of 10 ng/ml AHT during subculture for 3.5 h. The mean levels of adhesion and statistical significances were determined and expressed as described in Figure 1.

### Contribution of T1SS-secreted non-fimbrial adhesins to adhesion to lettuce

A small subpopulation of STM expresses SiiE under laboratory conditions (3.5 h subculture in LB). Enhanced surface expression of SiiE was achieved by AHT-induced expression of *hilD*, the central transcriptional activator of SPI1/SPI4 genes (31), as generation of a vector for Tet-on expression of the *sii* operon failed. The frequency of SiiE-expressing STM increased from approximately 12% under native conditions, to over 80% after AHT-induced *hilD* expression (28). The increased surface expression of SiiE in SR11 Δ12 correlated with significantly decreased adhesion to lettuce (Figure 3A), while a SPI4-deletion strain showed significantly increased adhesion. Since *hilD* expression also affects the expression of SPI1 and associated effector proteins, we tested the expression of *hilD* in strains harboring deletions in SPI4 or Δ*invC*, encoding the ATPase subunit of SPI1-T3SS. In both cases, neither the increased expression of SPI1-T3SS genes in STM ΔSPI4, nor the increased expression of SPI4 genes in Δ*invC* led to altered adhesion compared to background strain SR11 Δ12. Therefore, possibly the increased simultaneous expression of genes in SPI4 and SPI1 impaired adhesion to lettuce leaves.

**Figure 3:**
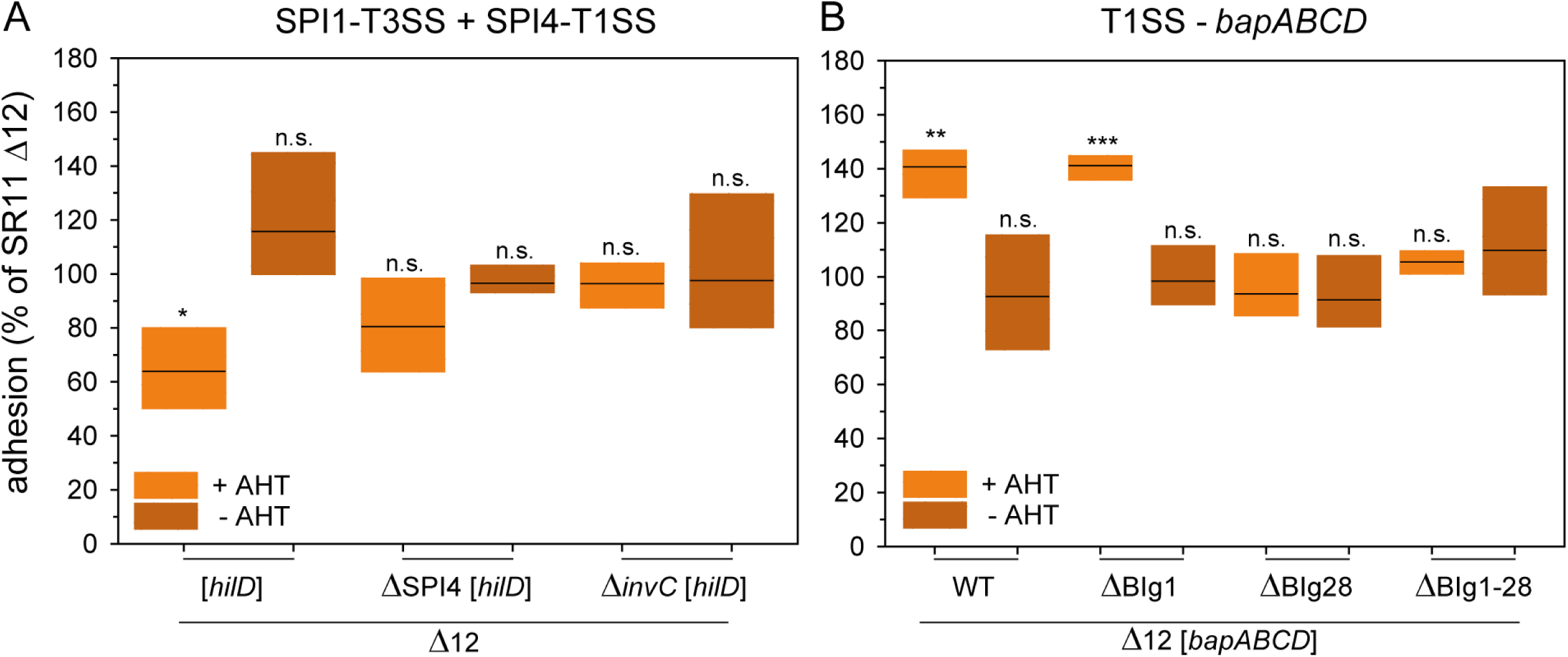
Impact of T1SS-secreted adhesins and *hilD* expression on adhesion of STM to lettuce. A) Lettuce grown under aseptic conditions was infected with STM strain SR11 Δ12 with overexpression of regulator *hilD* for analysis of the SPI4-encoded T1SS-secreted adhesin SiiE, and the SPI1-encoded T3SS. B) Adhesion of SR11 Δ12 stains with surface expression of T1SS-secreted adhesin BapA or truncated forms of BapA. Expression of adhesins was induced by addition of AHT if indicated. The mean levels of adhesion and statistical significances were determined as described in Figure 1.

AHT-induced expression of *bapABCD* led to significantly increased adhesion to lettuce, whereas deletion of *bapABCD* did not affect adhesion (Figure 3B). To achieve a better understanding of BapA binding to lettuce leaves, we used various truncated forms of BapA (28). Alleles of *bapA* with deletions of BIg1, BIg28, or BIg1-28 were expressed by AHT induction. Synthesis and secretion of mutant forms of BapA were confirmed by flow cytometry in prior work (28), and revealed that deletion of BIg1-28 ablated surface expression of BapA. The deletion of BIg1 showed a similar significantly increased adhesion as observed before for full-length BapA, indicating no contribution of BIg1 in adhesion to lettuce. Further, deletion of BIg28 resulted in adhesion comparable to background strain SR11 Δ12. Thus, deletion of only one BIg domain can be critical for enhanced adhesion of STM by BapA to lettuce. Deletion of BIg1-28 resulted in adhesion comparable to background strain SR11 Δ12, which is in line with loss of BapA surface expression.

### Contribution of autotransported adhesins to adhesion to lettuce

The AHT-induced expression of *shdA* led to a significantly decreased adhesion to lettuce leaves (75% mean, Figure 4A), as well as deletion of *shdA* (63% in average). AHT-induced expression of *sadA* did not alter adhesion to lettuce, but deletion of *sadA* resulted in significantly reduced adhesion (77% in average). The AHT-induced expression of *misL* revealed a significant increased adhesion to lettuce compared to background strain SR11 Δ12 (Figure 4B). To gain further insight into the contribution of MisL in binding to lettuce leaves, we decided to test truncated forms of MisL in adhesion to lettuce. MisL contains of a cleavable signal sequence (aa1-28) sec-dependent secretion into periplasm, a translocation domain (aa677-955) forming a β-barrel in the outer membrane and mediating transport of the passenger domain (aa29-676) across the outer membrane. The passenger domain is further divided into a non-conserved region and a conserved region compared to other, non-protease autotransported adhesins (24). C. W. Dorsey et al. (24) revealed that the non-conserved passenger domain (aa29-281) is responsible for binding of MisL to fibronectin, and with lower affinity to collagen IV. Our truncated forms of MisL included a partial deletion of the non-conserved portion of the passenger domain (aa29-227), and a larger deletion of the entire passenger domain (aa29-479). Synthesis and secretion of truncated forms of MisL was confirmed by flow cytometry (Figure S2AB). Furthermore, potential effects on bacterial autoaggregation were investigated by microscopy (Figure S2C) and revealed lack of bacterial autoaggregation with or without AHT induction. Expression of truncated *misL* Δ29-227 and *misL* Δ29-479 revealed no increased adhesion to lettuce (Figure 4B).

**Figure 4:**
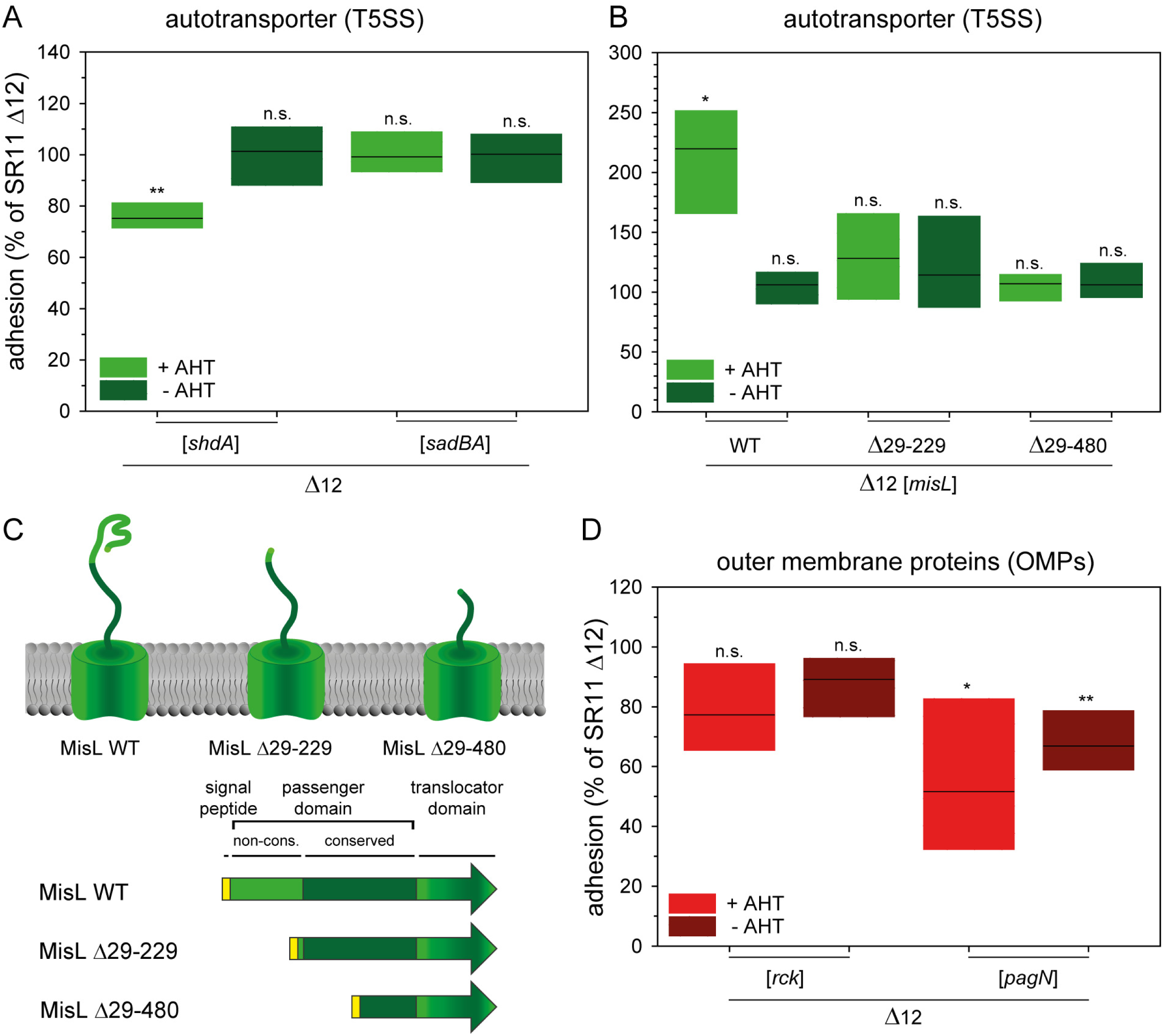
Impact of T5SS-secreted adhesins and outer membrane proteins on adhesion of STM to lettuce. Lettuce grown under aseptic conditions was infected with STM strain SR11 Δ12, or STM strain SR11 Δ12 with surface expression of T5SS-secreted adhesins ShdA or SadA (A), or MisL or truncated forms of MisL (B). Expression of the adhesins was induced by addition of AHT if indicated. C) Models for WT MisL and various truncated forms used in adhesion assays. D) For the analysis of outer membrane proteins, SR11 Δ12 strains expressing *rck* or *pagN* by induction of AHT were used. The mean levels of adhesion and statistical significances were determined as described in Figure 1.

In conclusion, the non-conserved region of MisL involved in binding to mammalian fibronectin and collagen IV is presumably also involved in the binding to lettuce leaves. Whether the impaired binding of truncated MisL is provoked by the fact that the fibronectin and/or collagen IV binding site is lost, or by the fact that truncated MisL has severely altered conformation has to be further investigated. Besides induced synthesis of MisL, deletion of *misL* also led to a significant increased adhesion to lettuce compared to background strain SR11 Δ12. However, STM Δ*misL* did not reach adhesion as observed for MisL synthesis.

### Contribution of OMP adhesins to adhesion to lettuce

In addition to SPI1-T3SS-mediated invasion of non-phagocytic cells by STM, two outer membrane proteins (OMPs) PagN and Rck were reported to mediate SPI1-T3SS-independent, zipper-like invasion (32, 33). Adhesive properties of these OMPs were proposed, which might also affect adhesion to non-mammalian organisms. AHT-induced expression of Rck, as well as deletion of *rck* resulted in no altered adhesion to lettuce leaves (Figure 4D). Expression of *pagN* led to a significantly reduced adhesion (52% mean), which was also observed for the non-induced samples (67% mean). However, Western blot analyses confirmed absence of PagN in non-induced cultures (23). In addition, a strain defective in *pagN* revealed a significantly reduced adhesion (43% mean).

### Contribution of flagella filaments and motility to adhesion to lettuce

Previous studies revealed contribution of flagella filaments and motility of STM in adhesion to various plant species (7, 34, 35). Here, we investigate the binding properties of the flagella filament, and the contribution of motility in the adhesion to lettuce leaves by using four distinct deletion strains. Loss of the flagella filaments (Δ*fliC* Δ*fljB*) resulted in significantly decreased adhesion to lettuce leaves (37% mean, Figure 5A). To bypass reduced interaction with leaf surfaces possibly resulting from loss of motility, contact was forced by centrifugation (5 min at 500 x g). However, centrifugation did not restore adhesion of STM to leaf surfaces (41% mean). To analyze the contribution of flagella filaments as putative adhesive structure, we used SR11 Δ12 Δ*motAB* defective in the energization of the flagella motor, but still harboring flagella filaments. This strain revealed significantly reduced adhesion in static and centrifuged samples (41% and 42% means, respectively). Therefore, presence of flagella filaments without rotation is not sufficient for adhesion to lettuce. Deletion of *cheZ*, resulting in tumbling only, also exhibited a significantly decreased adhesion to lettuce (static 45% and centrifuged 50% mean). A strain defective in *cheY*, restricted to smooth swimming, showed a slight but non-significantly decreased adhesion to lettuce leaves under static conditions (87% mean). After centrifugation, no altered adhesion (111% mean) was detected for Δ*cheY* compared to parental strain SR11 Δ12. Thus, we conclude a contribution of chemotactic motility of STM for adhesion to lettuce leaves.

**Figure 5:**
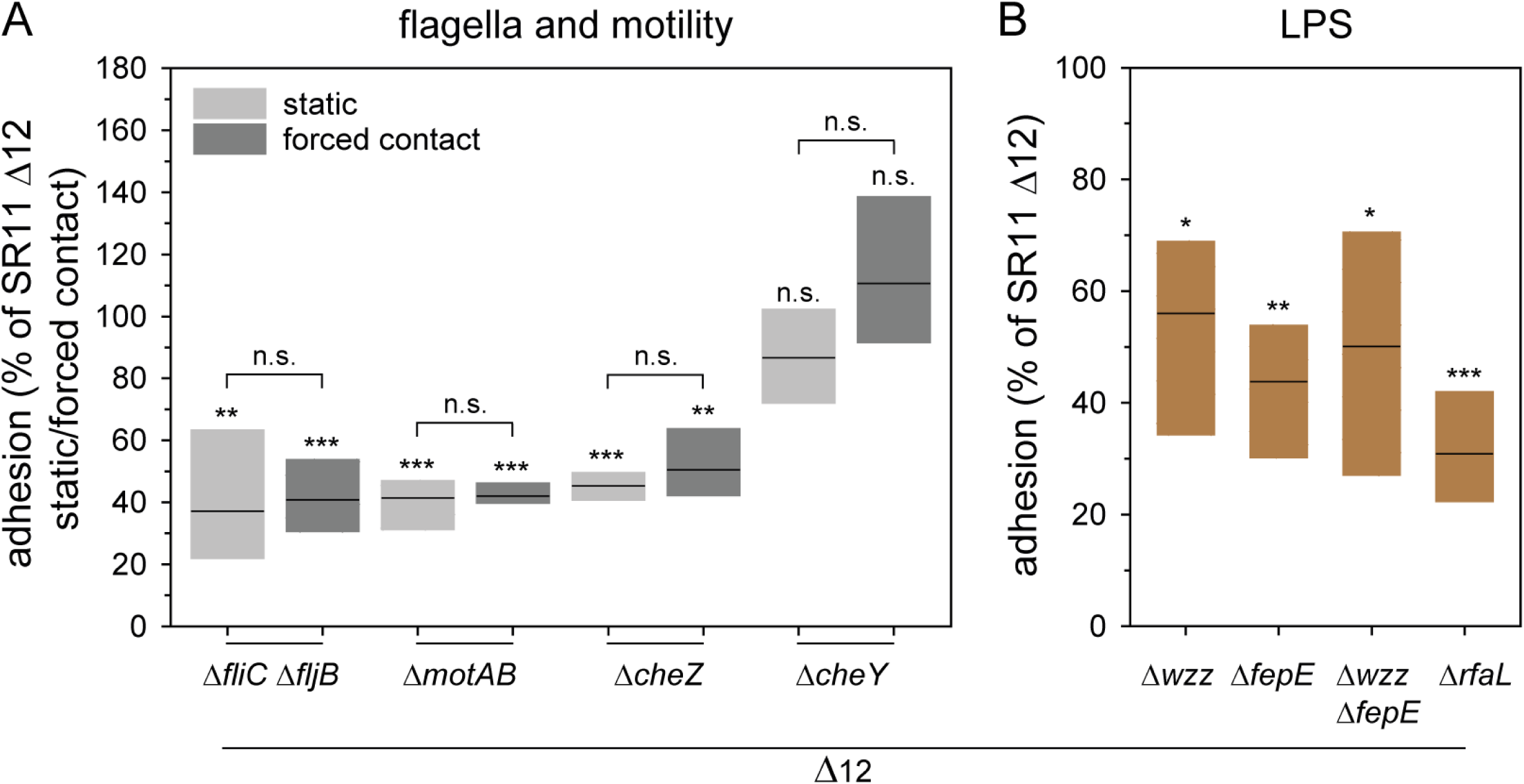
Impact of defects in motility or flagella assembly, or alteration in O-antigen length on STM adhesion to lettuce. A) Lettuce grown under aseptic conditions was infected with STM SR11 Δ12, or STM SR11 Δ12 with deletion of various genes required for motility or flagella-associated. B) Effect of mutations in genes resulting in lack (*rfaL*) or various truncations of O-antigen length (*wzz*_ST_, *fepE, wzz*_ST_ *fepE*). Infection was performed under static conditions (A, B), or by forced contact by centrifugation at 500 x g for 5 min to compensate effects of mutations in motility genes (A). The adhesion and the statistical significances were determined as described in Figure 1.

### Contribution of O-antigen to adhesion to lettuce

A main constituent of the bacterial cell envelope is LPS that stabilizes the cell envelope and protects Gram-negative bacteria against various environmental factors. Furthermore, LPS increases the negative charge of the cell envelope, and a putative adhesive role was reported (36). To analyze the contribution of LPS in adhesion to lettuce leaves, we used mutant strains lacking various genes involved in the biosynthesis and control of modal repeats of O-antigen (OAg) of LPS. In STM WT, a heterogeneous distribution of short chain OAg, long chain OAg (L-OAg), and very long chain OAg (VL-OAg) is found. Deletion of *wzz* results in the homogenous distribution of only VL-OAg, whereas in STM Δ*fepE* a homogenous distribution of only L-OAg is present. A strain defective in both genes (Δ*wzz* Δ*fepE*) only possesses short OAg (S-OAg). Deletion of *rfaL* results in LPS with core oligosaccharide (OS) lacking OAg. All deletion strains showed a significantly reduced adhesion to lettuce leaves compared to background strain SR11 Δ12 (Figure 5B, i.e. means of 56% for Δ*wzz*, 44% for Δ*fepE*, 31% for Δ*wzz* Δ*fepE*, and 50% for Δ*rfaL*). Hence, the heterogeneous distribution of L-OAg and VL-OAg on the bacterial surface is an important factor in the adhesion to lettuce leaves.

## Discussion

Here we addressed which factors of *S. enterica* are involved in adhesion to plant surfaces using a reductionist, synthetic approach with controlled surface expression of specific adhesive structures. All known adhesive structures encoded by STM were tested for their impact in adhesion to lettuce leaves. The results of this study are summarized in Figure 6.

**Figure 6:**
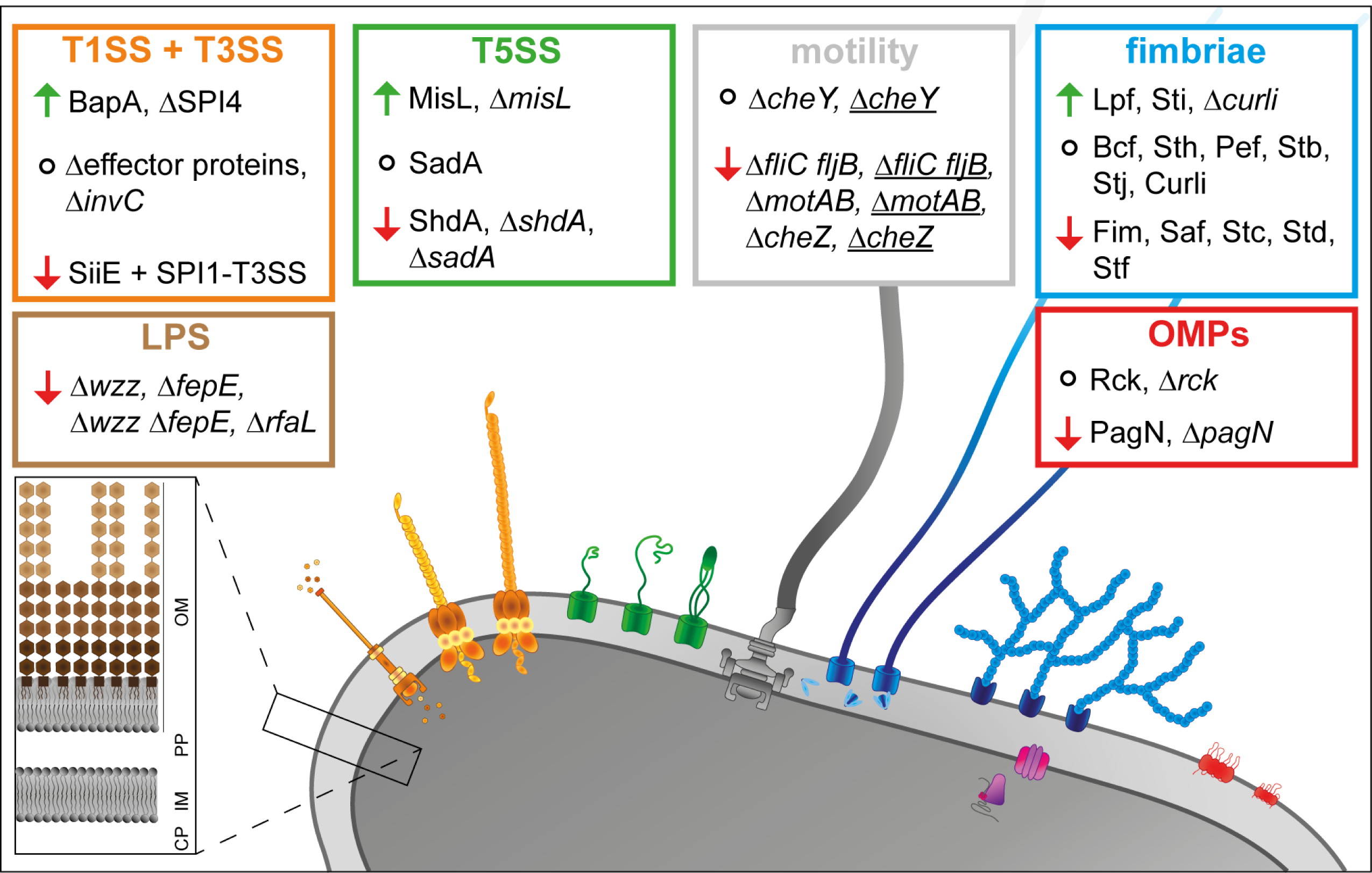
Overview of the role of analyzed factors in adhesion of STM to lettuce leaves. The absence of underlining indicates static samples, and underlining indicates centrifuged samples (forced contact). Arrows indicate increased (green) or decreased (red) adhesion, and circles indicate that adhesion was not altered. OM, outer membrane; PP, periplasm; IM, inner membrane; CP, cytoplasm.

The involvement of flagella of STM in adhesion to various salad species has been investigated before in several studies that reported decreased adhesion to basil, lettuce, and corn salad leaves for mutant strains lacking the flagella filament (7, 28, 37). These results are in line with our findings for decreased adhesion to lettuce leaves of an STM mutant strain lacking flagella filaments. Y. Rossez et al. (34) investigated the binding of pathogenic and nonpathogenic *E. coli* strains on a molecular level, revealing ionic binding of sulphated and phosphorylated plasma membrane lipids on *Arabidopsis thaliana* leaves. Whether the STM flagella filament also binds these molecules still has to be investigated. In addition to the flagella filament, Kroupitski *et al.* (7) found a contribution of chemotaxis, since a Δ*cheY* strain (only smooth swimming) showed decreased internalization in lettuce leaves. These authors hypothesized that lack of chemotaxis to higher sucrose concentrations close to stomata is affected, whereas the initial attachment of lettuce leaves was not affected. We investigated the involvement of motility in adhesion to lettuce leaves. A decreased adhesion was shown for strains deficient in energization and switching from CCW to CW flagella rotation (‘run only’), thus indicating the importance of directed motility and the capability of stopping on the leave surfaces. Bacteria only smooth swimming are probably not able to efficiently contact the leaf surface, or the initial adhesion is disrupted by the force generated by flagella rotating CCW. In summary, STM adhesion to lettuce leaves depends on motility, and in particular on the ability to switch flagella rotation from CCW (run) to CW (tumble).

The contribution of LPS of STM to virulence was investigated for mammalian cells in several studies before (38, 39). However, less is known about the involvement of STM LPS in adhesion to plants. In this study, we elucidated the importance of LPS with full length O-antigen for STM adhesion to lettuce leaves. The data indicate that LPS exhibits either specific binding properties for lettuce leaves or more probable, the surface charge and/or hydrophilicity of STM with intact LPS leads to an adhesion to lettuce leaves. This explanation for our finding of STM LPS involvement in adhesion to corn salad leaves is in line with a previous study (28). In studies on adhesion of pathogenic *E. coli*, lower attachment of LPS mutants to *A. thaliana* and romaine lettuce leaves was observed (40), indicating the importance of LPS in general for adhesion to plants. Further, M. H. de Moraes et al. (41) showed the involvement of STM LPS also for the persistence in tomatoes, revealing an impact in the entire infection process. However, changes of the bacterial cell surface charge due to altered LPS structure were not shown yet, but changes in biofilm formation were observed for bacterial plant pathogens with altered LPS structure. For example, altered LPS structure of *P. aeruginosa* caused decreased biofilm formation and affected virulence (42). Nevertheless, the involvement of LPS of STM in adhesion to plants, and effects of altered O-antigen structure on surface properties have to be further investigated.

In addition, the expression of fimbrial adhesins revealed increased adhesion to lettuce leaves if Sti or Lpf fimbriae were expressed. While expression was observed during infection of bovine ligated ileal loops (11), no specific ligands are knowns so far for Sti or Lpf. Whereas Sti fimbriae are absent in *S. enterica* serovar Typhi and in some strains of serovar Paratyphi A, Lpf are restricted to Typhimurium (43). A. J. Bäumler et al. (44) showed the involvement of Lpf fimbriae in adhesion to murine Peyer’s patches and to HEp-2 cells (44), and Lpf fimbriae contribute to long-term intestinal colonization of STM in resistant mice (45). Additionally, N. A. Ledeboer et al. (14) revealed 10-fold increased amounts of LpfE in biofilms, leading to the assumption that Lpf fimbriae are involved in biofilm formation during microcolony stage. Moreover, deletion of Lpf fimbriae resulted in loss of biofilm formation on chicken intestine. As for Lpf fimbriae, little is known about Sti fimbriae. A study by P. Laniewski et al. (46) revealed the involvement of Sti, Saf, Stc and curli fimbriae in murine infection model. Deletion of all four fimbrial operons resulted in a strain attenuated in a murine infection model, but if only one operon was deleted, no effect was observed. This indicates a possible redundancy of these fimbriae in murine infection, which were not observed for adhesion to lettuce leaves. Binding properties of Sti and Lpf fimbriae have to be further investigated, i.e. by glycoarrays (15).

Deletion of SPI4 encoding SiiE and its cognate T1SS led to an increased adhesion, whereas overexpression of SPI4 genes by overexpression of regulator *hilD* did not alter adhesion compared to background strain. These results indicate a contribution of SiiE for the whole bacterial population in STM adhesion to lettuce leaves. The same results were observed for adhesion to corn salad leaves (28).

We showed the involvement of BapA in adhesion to lettuce leaves. Whereas binding properties for specific glycostructures of BapA remain unclear, several studies revealed a contribution of BapA in biofilm formation (17, 22). Data obtained in this study after infection for 1 h were independent of biofilm formation by BapA. In our previous study, we revealed the involvement of BapA expression in adhesion to corn salad leaves leading to increased adhesion compared to background strain (28). Further, we tested the involvement of various BIg domains by using truncated forms of BapA and revealed a length-dependent adhesion of BapA to corn salad leaves, which was reduced by deletion of at least one BIg domain. Testing similar truncated forms of BapA in adhesion to lettuce leaves revealed suggested as similar role for BIg28, since binding mediated by BapA ΔBIg28 was reduced to the level of the background strain. In contrast, deletion of BIg1 did not affect adhesion compared to WT BapA. These observations indicate BIg-specific binding properties of BapA. However, the BapA topology has to be taken into consideration. While BIg28 is likely most distal to the outer membrane core, BIg1 is most proximal to the outer membrane and less likely to contact potential ligands. Moreover, *bapABCD* is highly conserved among *S. enterica* serovars (with some variations in *bapA*) indicating an important purpose during *S. enterica* lifestyle (47). Contribution of BapA encoded by other *S. enterica* serovars in adhesion to various salad species has to be tested in addition to the influence of longer infection times and involvement of BapA affecting biofilm formation on salad leaves. Here, we suggest to general role of STM BapA in adhesion to salad species.

In this study, we showed an involvement of autotransported MisL in adhesion to lettuce leaves. Expression of MisL is known to facilitates adhesion to CaCo2 and HeLa cells, enhancing biofilm formation (48), and contributing to intestinal colonization of mice (24). Furthermore, binding specificity of the non-conserved passenger domain (aa29-281) for fibronectin and, with lower affinity for collagen IV, were determined in extracellular matrix protein-binding assays (24). Deletion of the non-conserved transporter domain led to adhesion comparable to background strain indicating an involvement of fibronectin or collagen IV binding in adhesion to lettuce leaves. Purification of plant cell wall of *Pisum sativum* (peas) revealed a fibronectin-like protein (49), that may act as potentially ligands for MisL. However, direct contact to the plant cell wall for attaching bacteria is unlikely due to the barrier of overlaying epicuticular waxes. ShdA which is also known for fibronectin binding (50), showed no increased adhesion to lettuce leaves. Therefore, the ligand for MisL on lettuce leaves remains to be identified. The relevance of MisL in colonization of lettuce was previously investigated by Y. Kroupitski et al. (8). A genetic screen identified, together with *stfC* (encoding C/U fimbriae subunit) and *bcsA* (cellulose synthase subunit), *misL* as upregulated in STM on lettuce after storage for 7 days at 8 °C. Deletion of *misL* led to decreased survival on lettuce leaves under cold storage conditions. Interestingly, the effect of *misL* on survival was dependent on the presence of lettuce and absent after sole cold storage (8).

In this study, we showed for the first time the contribution of directed motility, an intact LPS layer, and expression of various adhesive structures of STM in adhesion to lettuce leaves. We revealed expression of Lpf or Sti fimbriae, T1SS-secreted BapA, or autotransported MisL leads to enhanced adhesion MisL to lettuce leaves. To gain further insight in the adhesion of STM to salad, BapA has to be further investigated possibly revealing common adhesive effects on plants in general. Furthermore, expression of all adhesive structures, especially adhesive structures involved in adhesion to salad, have to be further examined with regard to their native expression. For this purpose, we suggest transcriptomics or proteomics analyses of STM grown under various environmental conditions. In summary, with this work we further contributed to the understanding of the interaction of *S. enterica* adhesion to salad.

## Materials and Methods

### Bacterial strains and culture conditions

Bacterial strains used in this study are listed in Table 1. Unless otherwise stated, bacteria were grown in LB (lysogeny broth) medium, or on LB agar containing antibiotics for selection of specific markers if required to maintain plasmids listed in Table 2. Carbenicillin (Carb) and kanamycin (Km) were used to a final concentration of 50 µg/ml. For the induction of the Tet-on system, anhydrotetracycline (AHT) was added ad final concentrations of 10-100 ng/ml.

**Table 1.**
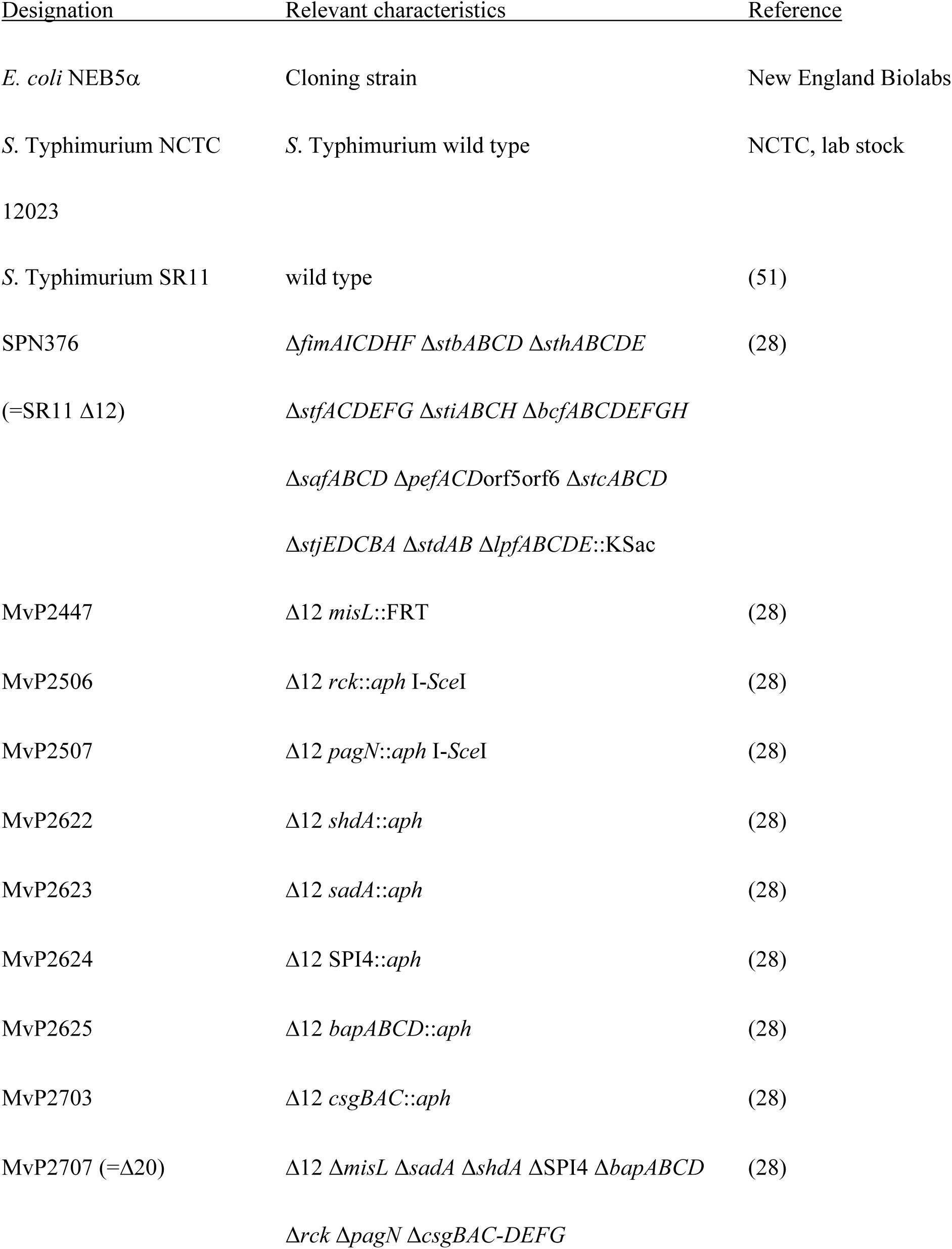
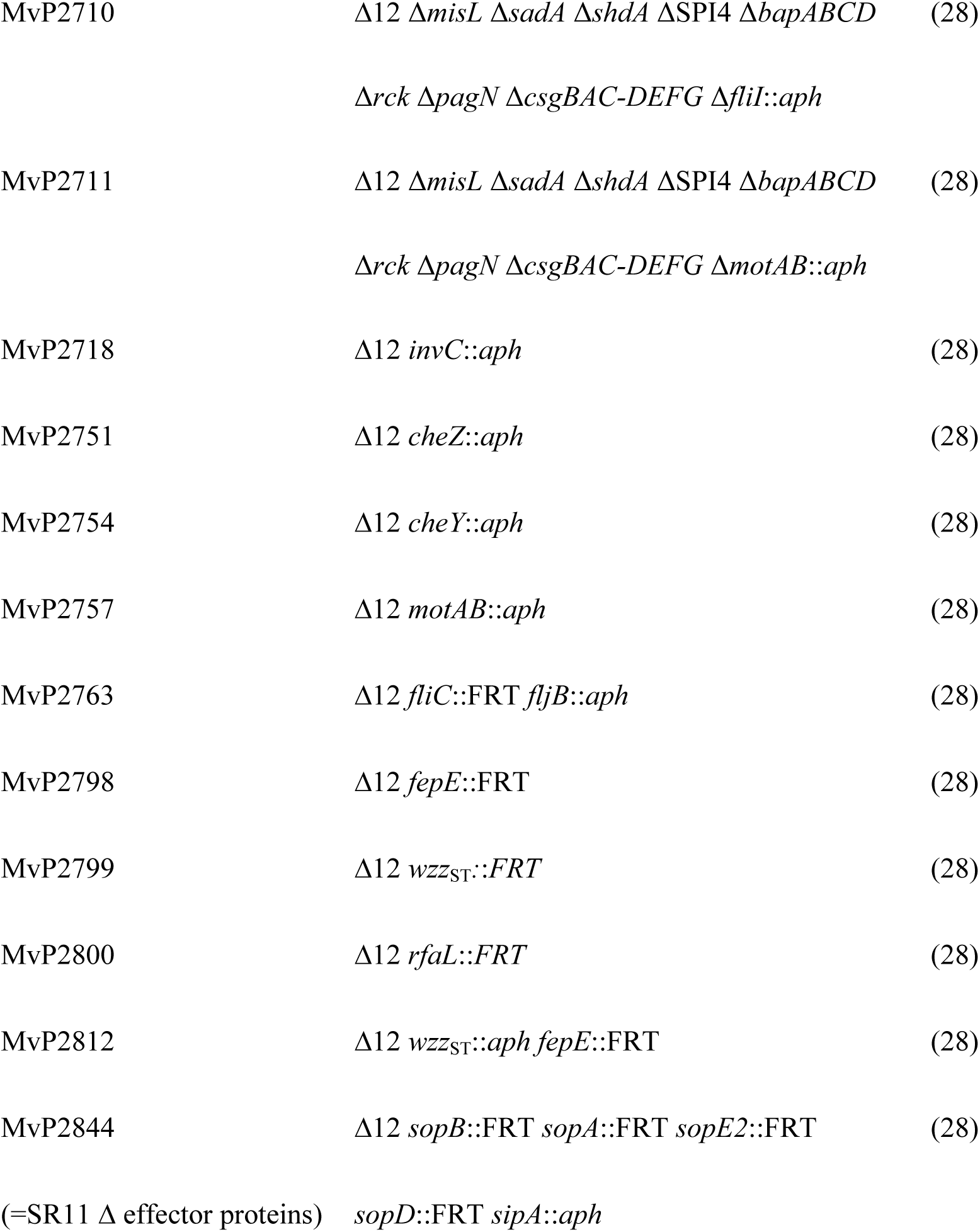
Bacterial strains used in this study.

**Table 2.**
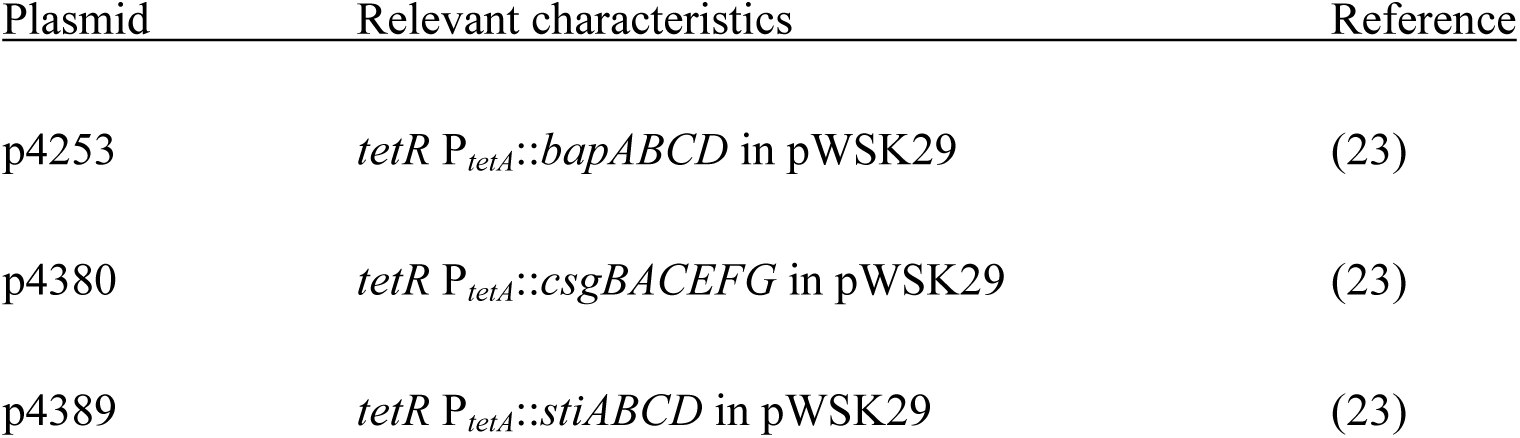
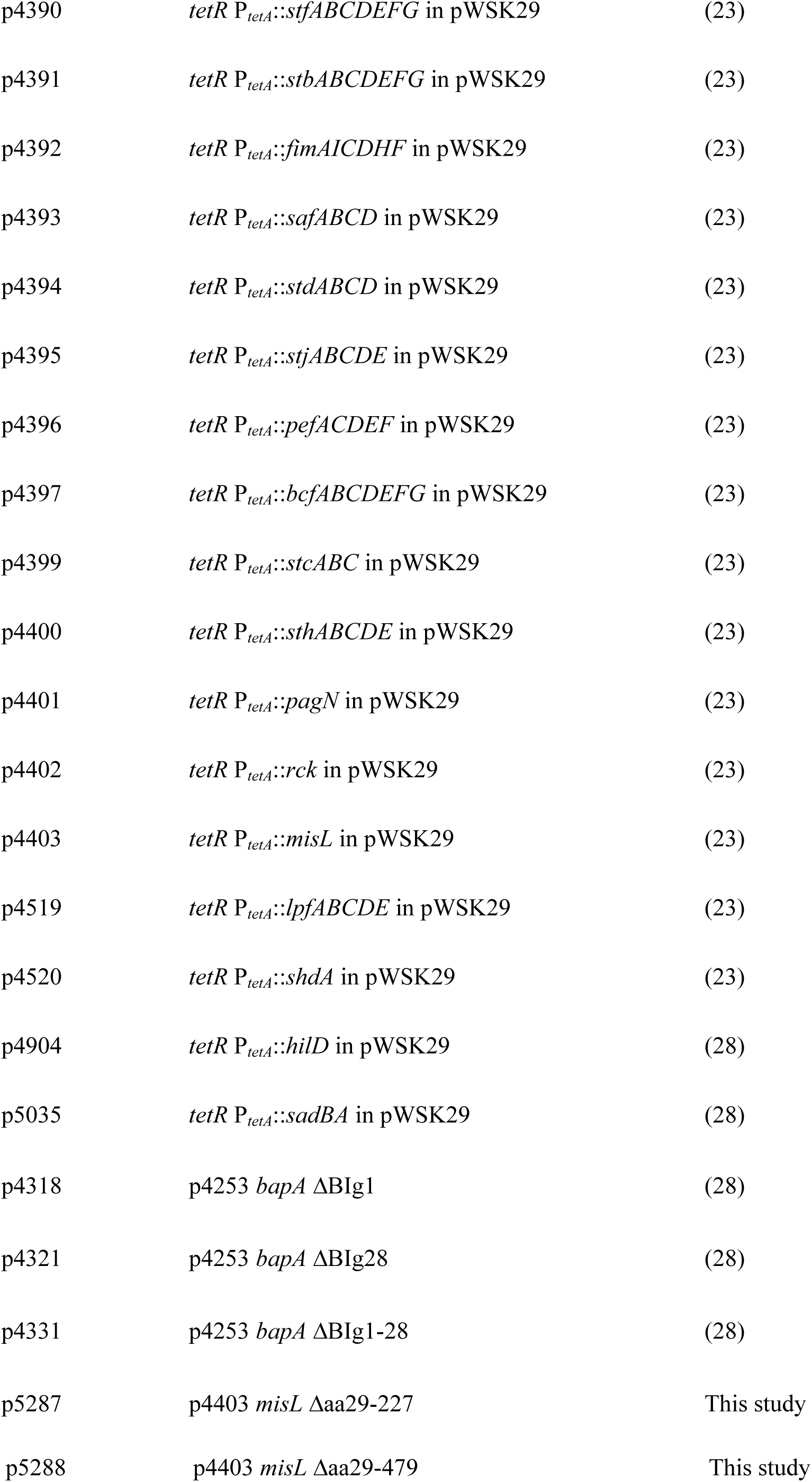
Plasmids used in this study.

**Table 3.**
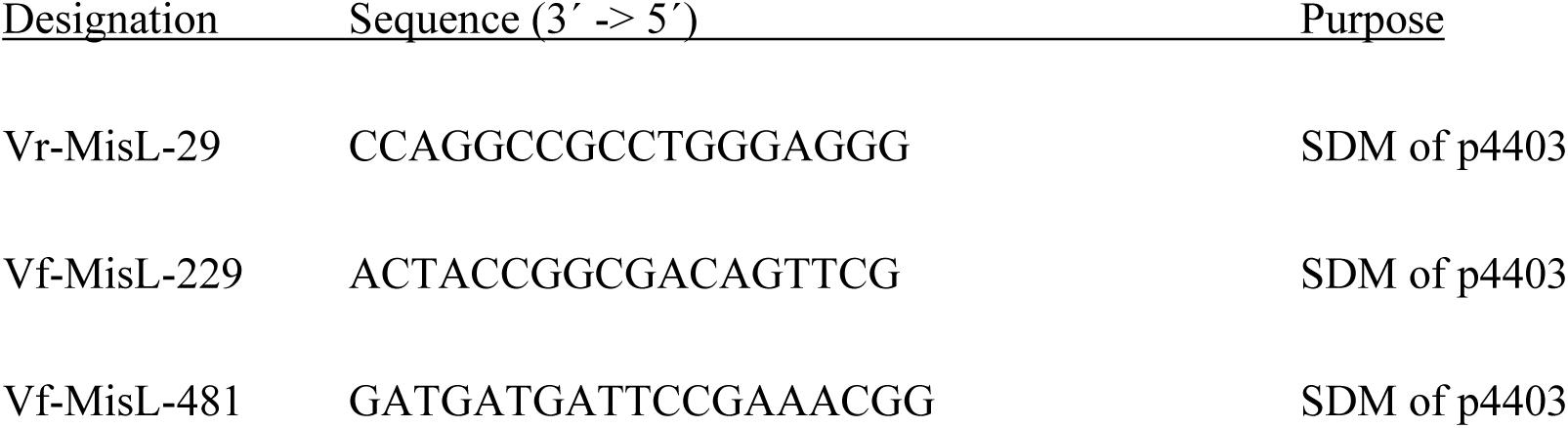
Oligonucleotides used in this study.

### Construction of strains and plasmids

For the construction of plasmids encoding various truncated forms of *misL* under the Tet-on system, plasmid p4403 was used as template. The plasmid was amplified by using oligonucleotides listed in Table 2, resulting in lack of various amounts of codons. Amplified products were reassembled by site-directed mutagenesis (SDM) kit according to manufacturer’s protocol (NEB). Sequence-confirmed plasmids were electroporated in STM SR11 Δ12.

### Cultivation of sterile grown lettuce and corn salad

Lettuce seeds (*Lactuca sativa* L. cultivar Tizian) and corn salad seeds (*Valerianella locusta* Verte à cour plein 2, N.L. Chrestensen Erfurter Samen-und Pflanzenzucht) were kindly provided by Dr. Adam Schikora and Dr. Sven Jechalke (Justus-Liebig University Giessen). Lettuce seeds were sterilized by 3% NaClO for 4 min thereby inverting the tube manually. Further, lettuce seeds were washed four times with sterile H_2_O_dd_ and directly planted on Murashige-Skoog (MS) agar (per liter: 2.2 g MS medium including vitamins, Duchefa Biochemie #M0222; 10 g agar; 5 g sucrose) in sterile plastic containers with air filter (round model 140 mm; Duchefa Biochemie, #E1674). Lettuce seeds were kept in the dark for 1 day at RT, and then further cultivated at 20 °C with a 12-h/12-h day-night-rhythm for 4 weeks. Corn salad seeds were sterilized by 70% EtOH for 1 min followed by 3% NaClO for 2 min. Seeds were washed thrice with sterile H_2_O_dd_ and dried for 30 min. Corn salad seeds were planted on MS agar (2.2 g MS medium including vitamins, Duchefa Biochemie #M0222; 10 g agar; 0.5 g MES per liter; pH 5.4) in sterile plastic containers with air filter as above at 20 °C with a 12 h/12 h day-night-rhythm for 8 weeks.

### Adhesion to lettuce and corn salad

Infection of lettuce with *Salmonella* was done as previously described in Elpers *et al.* (28) for infection of corn salad. In short, for each condition and strain three leaf discs of lettuce or corn salad were infected with STM strains diluted 1:31 from o/n cultures for 3.5 h. Infection was performed for 1 h at RT under static conditions, or for 55 min at RT after centrifugation of 5 min at 500 x g. After infection, leaf discs were washed with phosphate-buffered saline (PBS), homogenized and lysates were plated on MH (Mueller Hinton) agar plates, and incubated o/n at 37 °C. A non-infected sample was used in every assay to ensure sterility of the lettuce or corn salad plants used.

### Flow Cytometry

For analysis of surface expression of MisL by flow cytometry, approximately 6×10^8^ bacteria were washed in PBS and then fixed with 3% paraformaldehyde in PBS for 20 min at RT. Bacteria were blocked with 2% goat serum in PBS for 30 min and afterwards stained with the specific primary antiserum rabbit α MisL (1:1,000) o/n, 4 °C. Staining with secondary antibody goat α rabbit-Alexa488 (1:2,000) was performed for 1 h at RT. Bacteria were measured with a Attune NxT Flow Cytometer (Thermo Fisher) and analyzed using Attune NxT software, version 3.11.

### Autoaggregation analysis

For analysis of autoaggregation of strains expressing MisL and various truncated forms of MisL, subcultures used for the infection of lettuce were diluted to 1× 10^8^ bacteria/ml in PBS. 7 µl of bacterial suspension were imaged by Zeiss AxioObserver with brightfield microscopy with a 40x objective. Images were recorded by an AxioCam, and data were analyzed using ZEN 2012.

## Acknowledgements

This work was supported by the Bundesanstalt für Landwirtschaft und Ernährung (BLE) by projects PlantInfect and PlantInfect2, grant 2813HS027). Further support by the DFG by grant SFB 944, project Z is kindly acknowledged. We thank the members of the PlantInfect consortium for fruitful discussion and exchange of reagents. The systematic analyses of GEO data for adhesin expression, and critical comment on the manuscript by Janina Noster are gratefully acknowledged. We like to thank Andreas J. Bäumler (UC at Davis) for sharing antiserum against MisL.

